# Control of Vein-Forming, Striped Gene-Expression by Auxin Signaling

**DOI:** 10.1101/2020.09.28.317644

**Authors:** Anmol Krishna, Jason Gardiner, Tyler J. Donner, Enrico Scarpella

## Abstract

Activation of gene expression in striped domains is a key building block of biological patterning, from the recursive formation of veins in plant leaves to that of ribs and vertebrae in our bodies. In animals, gene expression is activated in striped domains by the differential affinity of broadly expressed transcription factors for their target genes and the combinatorial interaction between such target genes. In plants, how gene expression is activated in striped domains is instead unknown. We address this question for the broadly expressed MONOPTEROS (MP) transcription factor and its target gene *ARABIDOPSIS THALIANA HOMEOBOX FACTOR8* (*ATHB8*). We find that *ATHB8* promotes vein formation and that such vein-forming function depends on both levels of *ATHB8* expression and width of *ATHB8* expression domains. We further find that *ATHB8* expression is activated in striped domains by a combination of (1) activation of *ATHB8* expression through binding of peak levels of MP to a low-affinity MP-binding site in the *ATHB8* promoter and (2) repression of *ATHB8* expression by MP target genes of the *INDOLE-3-ACETIC-ACID-INDUCIBLE* family such as *BODENLOS*. Our findings suggest that a common regulatory logic controls activation of gene expression in striped domains in both plants and animals despite the independent evolution of their multicellularity.

## INTRODUCTION

Narrow stripes of gene expression are fundamental units of biological patterning (e.g., (Macdonald et al., 1986; Takahashi et al., 2007; Mallarino et al., 2016)). Therefore, how multicellular organisms activate gene expression in narrow stripes is a central question in biology. In animals, where this question has been investigated extensively, broadly expressed transcription factors activate expression of their target genes in narrow stripes by (1) differential affinity of such transcription factors for their binding sites in target genes and (2) combinatorial interactions between transcription-factor-encoding target genes (Ashe and Briscoe, 2006; Rogers and Schier, 2011; Hironaka and Morishita, 2012; Sagner and Briscoe, 2017). For example, the transcription factor Dorsal forms a ventral-to-dorsal gradient in Drosophila embryos (reviewed in (Reeves and Stathopoulos, 2009)). Expression of Dorsal target genes with high-affinity Dorsal-binding sites is activated already at low levels of Dorsal, whereas expression of Dorsal target genes with low-affinity Dorsal-binding sites is activated only at high levels of Dorsal. However, this mechanism alone is insufficient to account for the expression of Dorsal target genes in stripes: interaction between Dorsal target genes themselves is also required: Dorsal activates expression of *snail*, which encodes a transcription factor that represses the expression of the Dorsal target gene *ventral nervous system defective*. Thus, expression of some Dorsal target genes such as *ventral nervous system defective* is repressed at high levels of Dorsal, at which *snail* is expressed, but activated at lower levels of Dorsal, at which *snail* is not expressed.

In plants too, broadly expressed transcription factors activate expression of their target genes in narrow stripes (e.g., (Brady et al., 2011)); however, how those broadly expressed transcription factors do so is unclear. Here we addressed this question for the *MONOPTEROS* (*MP*) - *ARABIDOPSIS THALIANA HOMEOBOX8* (*ATHB8*) pair of Arabidopsis genes (Baima et al., 1995; Hardtke and Berleth, 1998). *ATHB8* expression is activated in single files of isodiametric ground cells of the leaf (Kang and Dengler, 2004; Scarpella et al., 2004). *ATHB8*-expressing ground cells will elongate into procambial cells — the precursors to all vascular cells — and are therefore referred to as preprocambial cells (Kang and Dengler, 2004; Scarpella et al., 2004; Sawchuk et al., 2007; Marcos and Berleth, 2014). Activation of *ATHB8* expression in narrow preprocambial stripes depends on binding of the broadly expressed MP transcription factor to a low-affinity MP-binding site in the *ATHB8* promoter (Donner et al., 2009). However, the biological relevance of activation of *ATHB8* expression by MP is unclear: whereas *MP* promotes vein formation (Przemeck et al., 1996), *ATHB8* seems to have only transient and conditional functions in vein network formation (Baima et al., 2001; Donner et al., 2009).

Here we show that *ATHB8* promotes vein formation and that both levels of *ATHB8* expression and width of *ATHB8* expression domains are relevant to vein formation. Finally, we show that *ATHB8* expression is restricted to narrow preprocambial stripes by a combination of (1) activation of *ATHB8* expression through binding of peak levels of MP to a low-affinity MP-binding site in the *ATHB8* promoter and (2) repression of *ATHB8* expression by MP target genes of the *INDOLE-3-ACETIC-ACID-INDUCIBLE* family.

## RESULTS & DISCUSSION

### Response of Vein Network Formation to Changes in ATHB8 Expression and Activity

To understand how in plants broadly expressed transcription factors activate expression of their target genes in narrow stripes, we chose the *MP* – *ATHB8* pair of Arabidopsis genes. During leaf development, the broadly expressed MP transcription factor directly activates *ATHB8* expression in narrow preprocambial stripes that mark the position where veins will form (Donner et al., 2009), but the biological relevance of the interaction between the two genes is unclear.

That *MP* promotes vein formation is known (Przemeck et al., 1996), but the function of *ATHB8* in this process is unresolved: *athb8* mutants seem to have only transient and conditional defects in vein network formation, and the mutants have normal vein patterns (Baima et al., 2001; Donner et al., 2009). Therefore, we first asked whether *ATHB8* had any permanent functions in vein network formation. To address this question, we characterized the vein networks in mature first leaves of the *athb8-11* and *-27* loss-of-function mutants (Prigge et al., 2005) (Table S1) — and of other genotypes in our study — by means of four descriptors: a cardinality index, a continuity index, and a connectivity index (Verna et al., 2015), and a cyclicity index.

The cardinality index is a proxy for the number of “veins” (i.e. stretches of vascular elements that contact other stretches of vascular elements at least at one of their two ends) in a network. The continuity index quantifies how close a vein network is to a network with the same pattern but in which at least one end of each “vein fragment” (i.e. a stretch of vascular elements that is free of contact with other stretches of vascular elements) contacts a vein. The connectivity index quantifies how close a vein network is to a network with the same pattern but in which both ends of each vein or vein fragment contact other veins. The cyclicity index is a proxy for the number of meshes in a vein network.

The cardinality index of both *athb8-11* and *-27* was lower than that of wild type (WT) (Fig. 1A–C,K), suggesting that *ATHB8* promotes vein formation.

**Figure 1.**
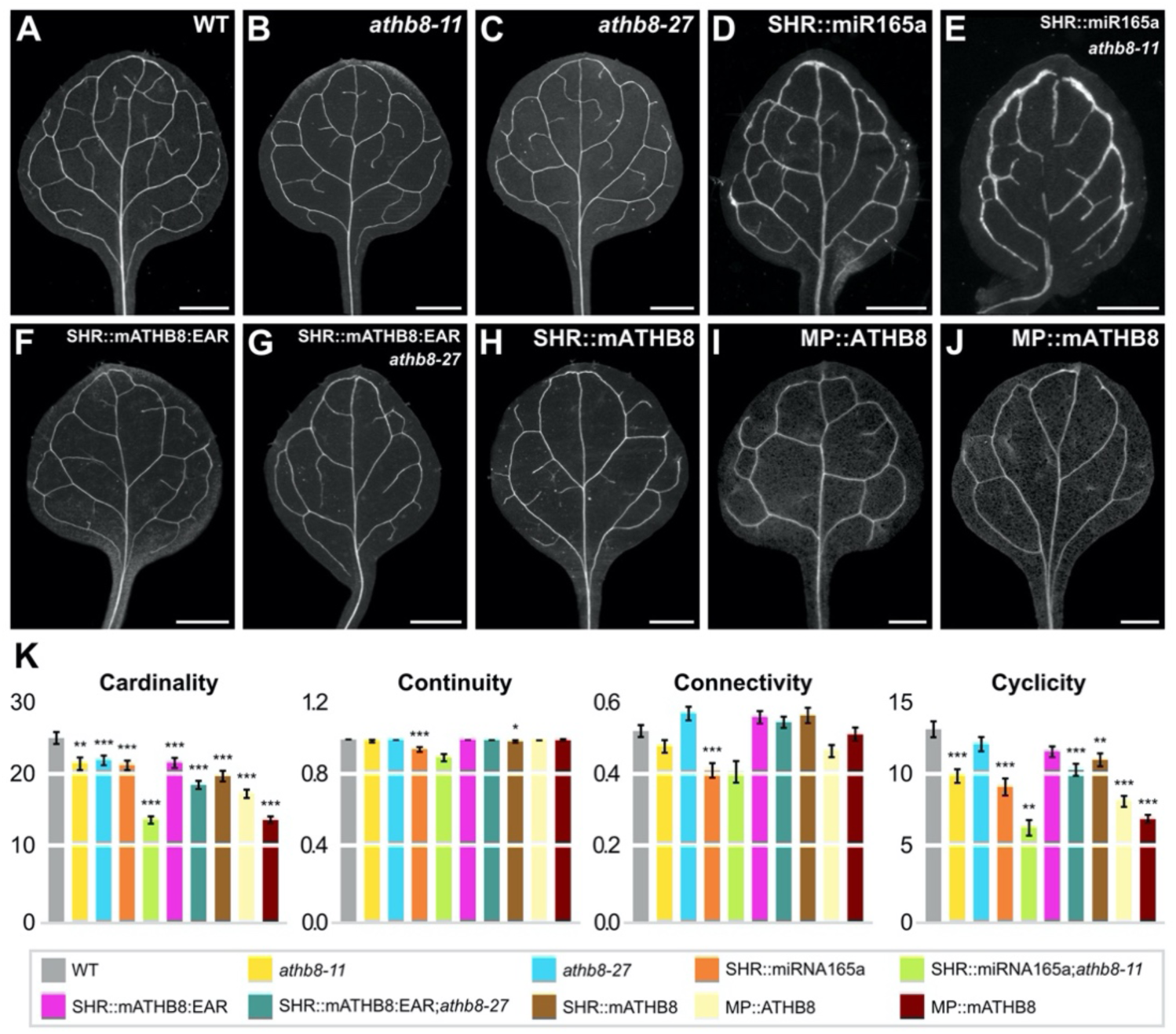
*ATHB8* Function in Vein Network Formation. (A–J) Dark-field illumination of cleared first leaves 14 days after germination (DAG); top right: genotype. (K) Cardinality, connectivity, and continuity index (mean ± SE) as defined in (Verna et al., 2015) and Materials & Methods; cyclicity index (mean ± SE) as defined in Materials & Methods. Difference between *athb8-11* and WT cardinality indices, between *athb8-27* and WT cardinality indices, between SHR::miR165a and WT cardinality indices, between SHR::miR165a;*athb8-11* and SHR::miR165a cardinality indices, between SHR::mATHB8:EAR and WT cardinality indices, between SHR::mATHB8:EAR;*athb8-27* and SHR::mATHB8:EAR cardinality indices, between SHR::mATHB8 and WT cardinality indices, between MP::ATHB8 and WT cardinality indices, between MP::mATHB8 and WT cardinality indices, between SHR::miR165a and WT continuity indices, between SHR::mATHB8 and WT continuity indices, between SHR::miR165a and WT connectivity indices, between *athb8-11* and WT cyclicity indices, between SHR::miR165a and WT cyclicity indices, between SHR::miR165a;*athb8-11* and SHR::miR165a cyclicity indices, between SHR::mATHB8:EAR;*athb8-27* and SHR::mATHB8:EAR cyclicity indices, between SHR::mATHB8 and WT cyclicity indices, between MP::ATHB8 and WT cyclicity indices, and between MP::mATHB8 and WT cyclicity indices was significant at *P*<0.05 (*), *P*<0.01 (**), or *P*<0.001 (***) by *F*-test and *t*-test with Bonferroni correction. Sample population sizes: WT, 58; *athb8-11*, 39; *athb8-27*, 32; SHR::miR165a, 51; SHR::miR165a;*athb8-11*, 64; SHR::mATHB8:EAR, 38; SHR::mATHB8:EAR;*athb8-27*, 28; SHR::mATHB8, 33; MP::ATHB8, 37; MP::mATHB8, 47. Scale bars: (A,I,J) 0.5 mm; (B,C,F,G,H) 1 mm; (D,E) 0.2 mm.

*ATHB8* encodes a transcription factor member of the HOMEODOMAIN-LEUCINE ZIPPER III (HD-ZIP III) family (Baima et al., 1995). To further test whether *ATHB8* promoted vein formation and to test whether *ATHB8* did so redundantly with other *HD-ZIP III* genes, we expressed *microRNA165a* (*miR165a*) — which targets all the *HD-ZIP III* genes (Zhou et al., 2007) — by the *SHORT-ROOT* (*SHR*) promoter — which drives expression in the *ATHB8* expression domain (Gardiner et al., 2011) — in both the WT and *athb8-11* backgrounds.

The cardinality index of SHR::miR165a was lower than that of WT and the cardinality index of SHR::miR165a;*athb8-11* was lower than that of SHR::miR165a (Fig. 1D,E,K), supporting that *ATHB8* promotes vein formation and suggesting that *ATHB8* does so redundantly with other *HD-ZIP III* genes.

HD-ZIP III proteins bind DNA as homo- or hetero-dimers (Sessa et al., 1998; Merelo et al., 2016). Therefore, to further test whether *ATHB8* promoted vein formation and whether *ATHB8* did so redundantly with other *HD-ZIP III* genes, we generated a dominant-negative version of the ATHB8 transcriptional activator (Baima et al., 2014) by fusing the *ATHB8* ORF to the sequence encoding the EAR (ETHYLENE-RESPONSIVE ELEMENT-BINDING PROTEIN-ASSOCIATED AMPHIPHILIC REPRESSION) portable repressor domain (Hiratsu et al., 2003). In the resulting ATHB8:EAR, we introduced silent mutations that abolish *miR165a*-mediated downregulation (Ohashi-Ito et al., 2013). We expressed the resulting mATHB8:EAR by the SHR promoter in both the WT and *athb8-27* backgrounds.

The cardinality index of SHR::mATHB8:EAR was lower than that of WT, and the cardinality index of SHR::mATHB8:EAR;*athb8-27* was lower than that of SHR::mATHB8:EAR (Fig. 1F,G,K), supporting that *ATHB8* promotes vein formation and that *ATHB8* does so redundantly with other *HD-ZIP III* genes.

We next asked whether levels of *ATHB8* expression and width of *ATHB8* expression domains were relevant to vein formation. To address this question, we used SHR::mATHB8, which overexpresses *ATHB8* in its expression domain; MP::ATHB8, which expresses *ATHB8* in the broader *MP*-expression domain; and MP::mATHB8, which overexpresses *ATHB8* in the *MP* expression domain.

The cardinality index of SHR::mATHB8 was lower than that of WT; the cardinality index of MP::ATHB8 was lower than that of SHR::mATHB8; and the cardinality index of MP::mATHB8 was lower than that of MP::ATHB8 (Fig. 1H–K). These results suggest that both levels of *ATHB8* expression and width of *ATHB8* expression domains are relevant to vein formation.

### Relation Between *ATHB8* Expression Domains and MP Expression Levels

Width of *ATHB8* expression domains is relevant to vein formation (Figure 1). Therefore, we asked how *ATHB8* expression is activated in narrow preprocambial stripes by the broadly expressed MP. We hypothesized that *ATHB8* preprocambial expression is activated in narrow stripes by binding of peak levels of the broadly expressed MP to a low affinity site in the *ATHB8* promoter. This hypothesis predicts that narrow stripes of *ATHB8* preprocambial expression correspond to peak levels of MP expression. To test this prediction, we simultaneously imaged expression of ATHB8::nCFP (nuclear CFP expressed by the *ATHB8* promoter) (Sawchuk et al., 2007) and MP::MP:YFP (MP:YFP fusion protein expressed by the *MP* promoter) in first leaves of the strong *mp-B4149* mutant (Weijers et al., 2005), whose defects were rescued by MP::MP:YFP expression (Fig. S1A– C) (Table S1).

*ATHB8* preprocambial expression can be reproducibly observed in midvein, first loops of veins (“first loops”), and second loops of first leaves, respectively 2, 3, and 4 days after germination (DAG) (Donner et al., 2009; Gardiner et al., 2011; Donner and Scarpella, 2013). At these stages, MP::MP:YFP was expressed in ATHB8::nCFP-expressing cells at higher levels than in cells flanking ATHB8::nCFP-expressing cells (Figure 2; Fig. S2A,B).

**Figure 2.**
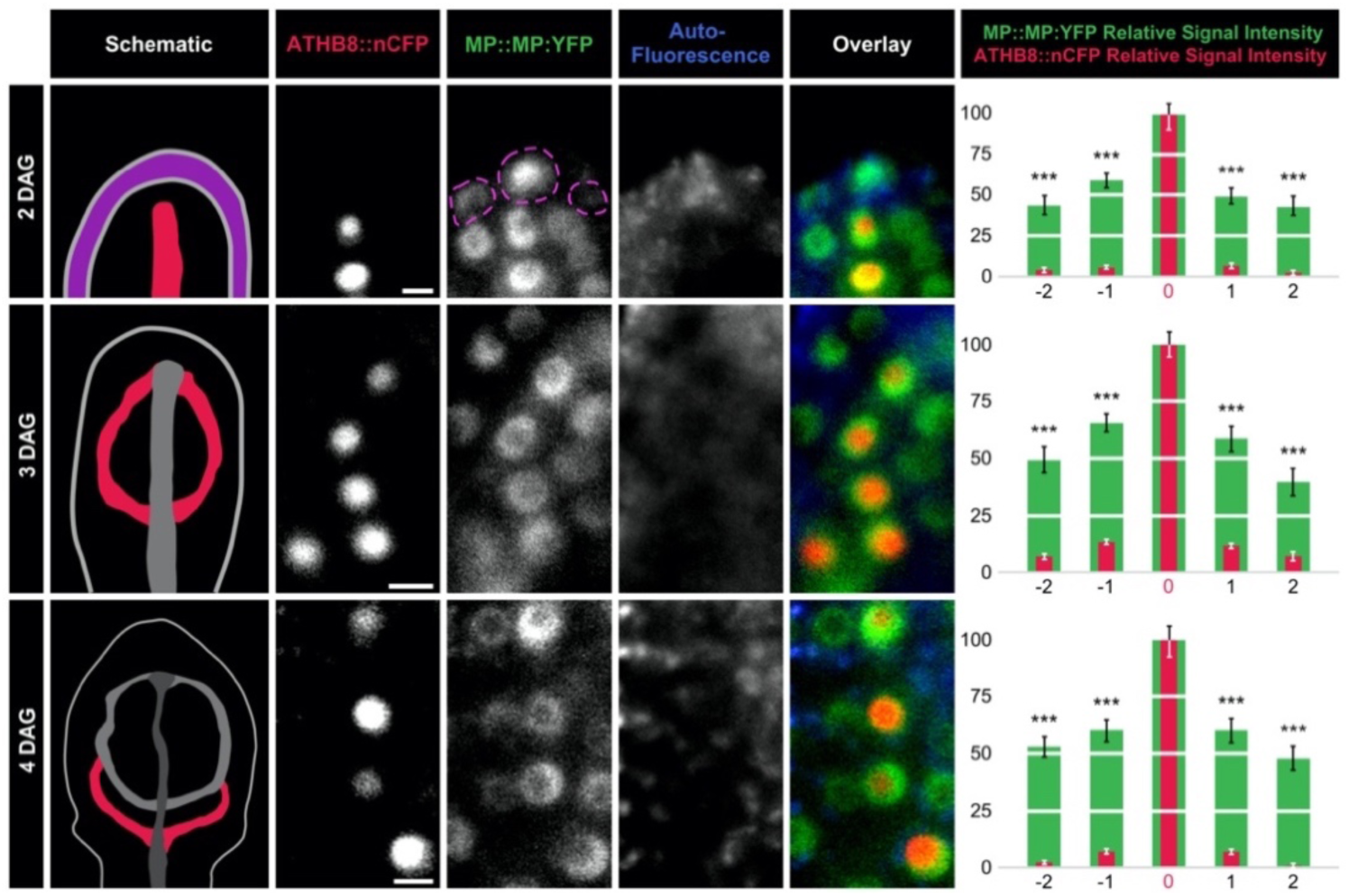
*ATHB8* and MP Expression Domains and Levels in Leaf Development. First leaves 2, 3, and 4 DAG. Column 1: schematics of leaves — imaged in columns 2–5 — illustrating onset of *ATHB8* expression (red) — imaged in column 2 - associated with formation of midvein (2 DAG), first loop (3 DAG), or second loop (4 DAG) (Donner et al., 2009; Gardiner et al., 2011; Donner and Scarpella, 2013); magenta: epidermis; increasingly darker gray: progressively older *ATHB8* expression domains. Columns 2–5: confocal laser scanning microscopy. Column 2: ATHB8::nCFP expression. Column 3: MP::MP:YFP expression; dashed magenta outline: MP::MP:YFP-expressing epidermal nuclei. Column 4: autofluorescence. Column 5: overlays of images in columns 2–4; red: ATHB8::nCFP expression; green: MP::MP:YFP expression; blue: autofluorescence. Column 6: MP::MP:YFP and ATHB8::nCFP expression levels (mean ± SE) in nuclei flanking ATHB8::nCFP-expressing nuclei (positions “-2”, “-1”, “1”, and “2”) relative to MP::MP:YFP and ATHB8::nCFP expression levels in nuclei co-expressing ATHB8::nCFP (position “o”) during formation of midvein (top), first loop (middle), or second loop (bottom). Difference between MP::MP:YFP expression levels in nuclei at position −2, −1, 1, or 2 and MP::MP:YFP expression levels in nuclei at position 0, and between ATHB8::nCFP expression levels in nuclei at position −2, −1, 1, or 2 and ATHB8::nCFP expression levels in nuclei at position 0 was significant at *P*<0.001 (***) by One-Way ANOVA and Tukey’s Pairwise test. MP::MP:YFP sample population sizes: 35 (2 DAG), 33 (3 DAG), or 32 (4 DAG) leaves; position −2: 30 (2 DAG), 45 (3 DAG), or 50 (4 DAG) nuclei; position −1: 63 (2 DAG), 72 (3 DAG), or 67 (4 DAG) nuclei; position 0: 70 (2 DAG), 75 (3 DAG), or 70 (4 DAG) nuclei; position 1: 58 (2 DAG), 47 (3 DAG), or 59 (4 DAG) nuclei; position 2: 24 (2 DAG), 19 (3 DAG), or 38 (4 DAG) nuclei. ATHB8::nCFP sample population sizes: 35 (2 DAG), 33 (3 DAG), or 32 (4 DAG) leaves; position −2: 29 (2 DAG), 43 (3 DAG), or 49 (4 DAG) nuclei; position −1: 57 (2 DAG), 70 (3 DAG), or 66 (4 DAG) nuclei; position 0: 63 (2 DAG), 73 (3 DAG), or 69 (4 DAG) nuclei; position 1: 52 (2 DAG), 46 (3 DAG), or 58 (4 DAG) nuclei; position 2: 23 (2 DAG), 19 (3 DAG), or 37 (4 DAG) nuclei. Scale bars (shown, for simplicity, only in column 2): 5 µm.

To test whether the differential expression of MP::MP:YFP in ATHB8::nCFP-expressing cells and in cells flanking ATHB8::nCFP-expressing cells were an imaging artifact, we compared expression levels of nCFP driven by a ubiquitously active promoter (RIBO::nCFP) (Gordon et al., 2007) in cells expressing ATHB8::nYFP (Sawchuk et al., 2007) and in cells flanking ATHB8::nYFP-expressing cells. We focused our analysis on second loops of 4-DAG first leaves, in which *ATHB8* preprocambial expression can be reproducibly observed (Donner et al., 2009; Gardiner et al., 2011; Donner and Scarpella, 2013).

Because levels of RIBO::nCFP expression in ATHB8::nYFP-expressing cells were no higher than those in cells flanking ATHB8::nYFP-expressing cells (Fig. S2D,E; Figure S3), we conclude that the differential expression of MP::MP:YFP in ATHB8::nCFP-expressing cells and in cells flanking ATHB8::nCFP-expressing cells is not an imaging artifact, and therefore that narrow stripes of *ATHB8* preprocambial expression correspond to peak levels of MP expression.

### Response of *ATHB8* Expression and Vein Network Formation to Changes in MP Expression

The hypothesis — that *ATHB8* preprocambial expression is restricted to narrow stripes by binding of peak levels of the broadly expressed MP transcription factor to a low affinity site in the *ATHB8* promoter — predicts that loss of *MP* function will lead to extremely weak, or altogether absent, *ATHB8* preprocambial expression, otherwise normally visible in second loops of 4-DAG first leaves (Donner et al., 2009; Gardiner et al., 2011; Donner and Scarpella, 2013). To test this prediction, we quantified ATHB8::nYFP expression levels in second loops of 4-DAG first leaves of the strong *mp-U55* mutant (Mayer et al., 1993; Donner et al., 2009).

Consistent with previous observations (Donner et al., 2009), ATHB8::nYFP expression levels were greatly reduced in *mp-U55*, leading to near-complete loss of ATHB8::nYFP preprocambial expression (Fig. 3A,B,F). Moreover, consistent with previous observations (Przemeck et al., 1996; Donner et al., 2009), near-complete loss of *ATHB8* preprocambial expression in *mp-U55* developing leaves was associated with networks of fewer meshes and fewer, less frequently continuous, and less frequently connected veins in *mp-U55* mature leaves (Fig. 3G,H,K).

**Figure 3.**
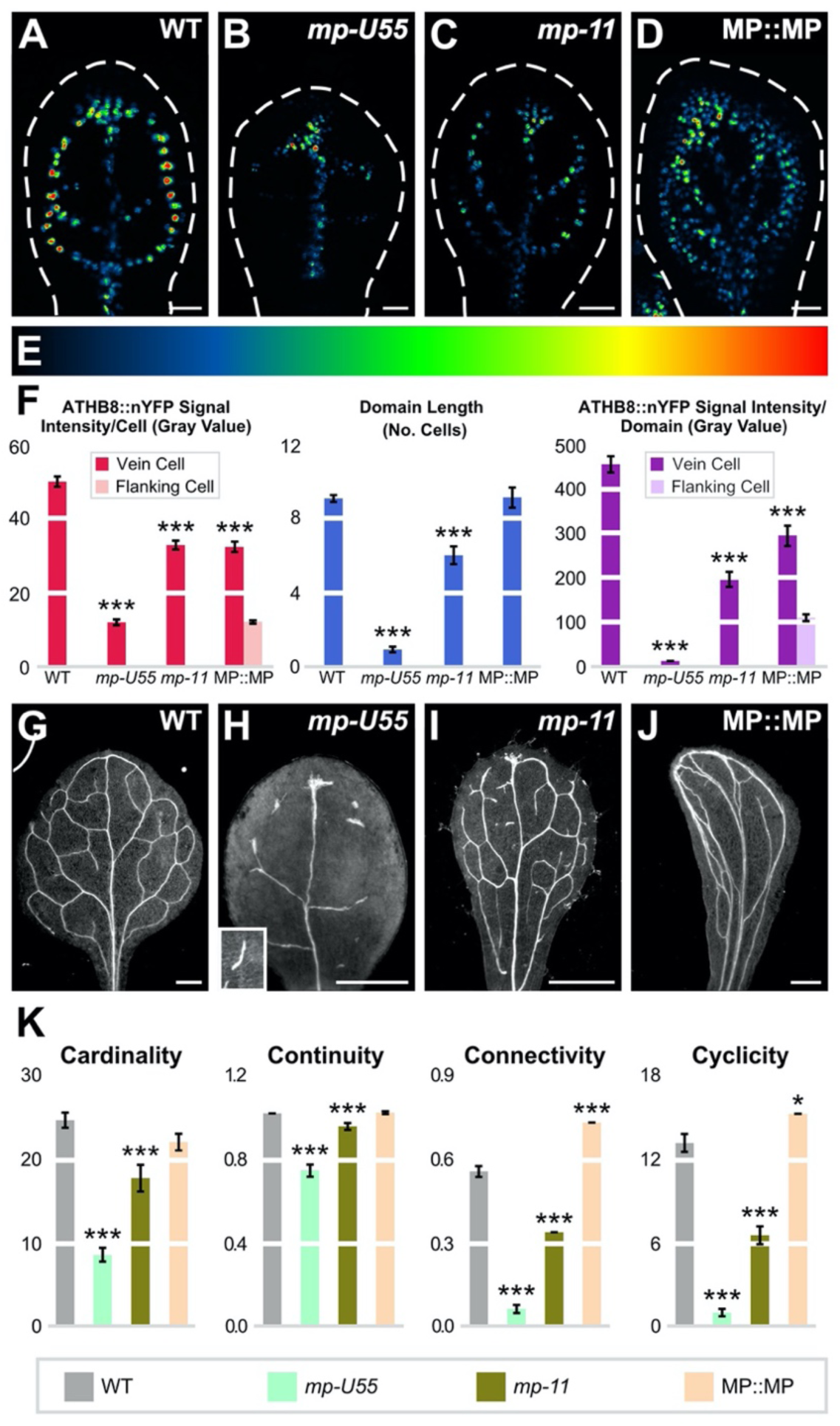
*MP* Expression, *ATHB8* Expression Domains and Levels, and Vein Network Formation. (A–D,G–J) Top right: genotype. (A–D) First leaves 4 DAG; confocal laser scanning microscopy; dashed white line: leaf outline; ATHB8::nYFP expression (look-up table — ramp in E — visualizes expression levels). (F) ATHB8::nYFP expression level per cell expressed as mean gray value ± SE, ATHB8::nYFP expression domain length expressed as mean number of cells ± SE, and ATHB8::nYFP expression levels per domain expressed as mean gray value ± SE. Difference between *mp-U55* and WT, between *mp-11* and WT, and between MP::MP and WT was significant at *P*<0.001 (***) by *F*-test and *t*-test with Bonferroni correction. Sample population sizes: 25 (WT), 72 (*mp-U55*), 27 (*mp-11*), or 24 (MP::MP) leaves; 345 (WT), 128 (*mp-U55*), 325 (*mp-11*), or 219 (MP::MP) vein cell nuclei, and 513 (MP::MP) flanking cell nuclei. (G-J) Dark-field illumination of cleared first leaves 14 DAG. (K) Cardinality index, connectivity index, and continuity index (mean ± SE) as defined in (Verna et al., 2015) and Materials & Methods; cyclicity index (mean ± SE) as defined in Materials & Methods. Difference between *mp-U55* and WT cardinality indices, between *mp-11* and WT cardinality indices, between *mp-U55* and WT continuity indices, between *mp-11* and WT continuity indices, between *mp-U55* and WT connectivity indices, between *mp-11* and WT connectivity indices, between MP::MP and WT connectivity indices, between *mp-U55* and WT cyclicity indices, between *mp-11* and WT cyclicity indices, and between MP::MP and WT cyclicity indices was significant at *P*<0.05 (*) or *P*<0.001 (***) by *F*-test and *t*-test with Bonferroni correction. Sample population sizes: WT, 39; *mp-U55*, 59; *mp-11*, 44; MP::MP, 41. Scale bars: (A-D) 25 µm; (G-J) 0.5 mm.

The hypothesis further predicts that lower levels of *MP* expression will lead to lower levels of *ATHB8* preprocambial expression. To test this prediction, we quantified ATHB8::nYFP expression levels in second loops of 4-DAG first leaves of the weak *mp-11* mutant, in which an insertion in the *MP* promoter (Odat et al., 2014) leads to ∼85% reduction in levels of WT *MP* transcript (Figure S4).

In *mp-11*, ATHB8::nYFP expression levels were lower and expression along the domain was more heterogeneous than in WT, leading to seemingly fragmented domains of weak ATHB8::nYFP preprocambial expression (Fig. 3A,C,F). Moreover, like in *mp-U55*, defects in *ATHB8* expression in *mp-11* developing leaves were associated with networks of fewer meshes and fewer, less frequently continuous, and less frequently connected veins in *mp-11* mature leaves (Fig. 3G,I,K). However, the vein network and *ATHB8* expression defects of *mp-11* were weaker than those of *mp-U55* (Fig. 3A–C,G–I,K).

The hypothesis also predicts that higher levels of the broadly expressed MP will lead to higher levels of *ATHB8* preprocambial expression in both vein and flanking cells, leading to broader *ATHB8* expression domains. To test this prediction, we overexpressed *MP* by its own promoter (MP::MP) — which led to ∼10-fold increase in *MP* expression levels (Figure S4) and which rescued defects of the strong *mp-B4149* mutant (Fig. S1A,B,D) (Table S1) — and quantified ATHB8::nYFP expression levels in second loops of 4-DAG MP::MP first leaves.

In MP::MP, ATHB8::nYFP expression levels were higher in flanking cells, leading to broad bands of ATHB8::nYFP expression; however, ATHB8::nYFP expression levels were lower in vein cells (Fig. 3A,D,F). Nevertheless, broad bands of *ATHB8* expression in MP::MP developing leaves were associated with abnormal vein networks in MP::MP mature leaves: veins ran close to one another for varying stretches of the narrow leaf laminae, then diverged, and either ran close to other veins or converged back to give rise to elongated meshes (Fig. 3G,J,K).

In summary, lower levels of *MP* expression lead to fragmented domains of *ATHB8* preprocambial expression, and loss of *MP* function leads to near-complete loss of *ATHB8* preprocambial expression. These observations are consistent with the hypothesis and suggest that *MP* expression levels below a minimum threshold are unable to activate *ATHB8* preprocambial expression. However, that higher levels of *MP* expression fail to lead to higher levels of *ATHB8* preprocambial expression in vein cells is inconsistent with the hypothesis and suggests that *MP* expression levels above a maximum threshold both activate and repress *ATHB8* preprocambial expression. These observations are unaccounted for by the hypothesis; therefore, the hypothesis must be revised.

### Response of *ATHB8* Expression and Vein Network Formation to Changes in MP Activity

*MP* expression levels above a maximum threshold both activate and repress *ATHB8* preprocambial expression (Figure 3). Activation of *ATHB8* preprocambial expression by MP is direct (Donner et al., 2009), but repression of *ATHB8* preprocambial expression by MP need not be: Repression of *ATHB8* preprocambial expression by MP could be mediated by BODENLOS (BDL)/INDOLE-3-ACETIC-ACID-INDUCIBLE12 (IAA12) (BDL hereafter), whose expression is activated by MP and which binds to MP and inhibits its transcriptional activity (Hamann et al., 2002; Hardtke et al., 2004; Weijers et al., 2005; Lau et al., 2011). Were repression of *ATHB8* preprocambial expression by MP mediated by BDL, *ATHB8* preprocambial expression would be reduced in the *bdl* mutant, in which the unstable BDL protein is stabilized (Dharmasiri et al., 2005). To test this prediction, we quantified ATHB8::nYFP expression levels in second loops of 4-DAG first leaves of the *bdl* mutant.

Like in *mp*, in *bdl* ATHB8::nYFP expression levels were lower and expression along the domain was more heterogeneous than in WT, leading to seemingly fragmented domains of weak ATHB8::nYFP preprocambial expression (Fig. 3A–C,F; Fig. 4A,B,G). Moreover, like in *mp*, defects in *ATHB8* expression in *bdl* developing leaves were associated with networks of fewer meshes and fewer, less frequently continuous, and less frequently connected veins in *bdl* mature leaves (Fig. 3G–I,K; Fig. 4H,I,M).

**Figure 4.**
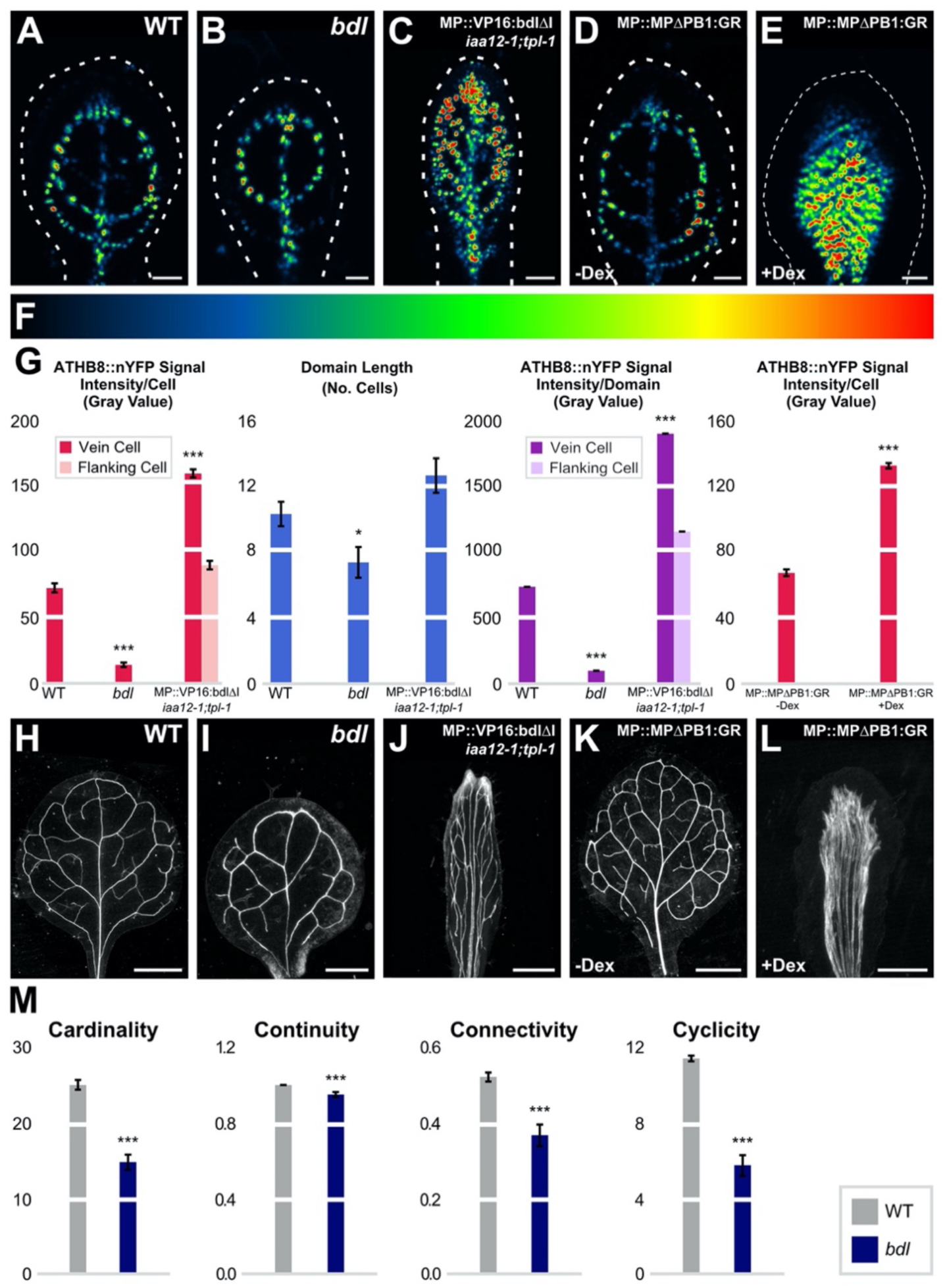
MP Activity, *ATHB8* Expression Domains and Levels, and Vein Network Formation. (A–E,H–L) Top right: genotype. (A–E) First leaves 4 DAG; confocal laser scanning microscopy; dashed white line: leaf outline; ATHB8::nYFP expression (look-up table — ramp in F — visualizes expression levels). (G) ATHB8::nYFP expression level per cell expressed as mean gray value ± SE, ATHB8::nYFP expression domain length expressed as mean number of cells ± SE, and ATHB8::nYFP expression levels per domain expressed as mean gray value ± SE. Difference between *bdl* and WT, between MP::VP16:bdlΔI;*iaa12-1*;*tpl-1* and WT, and between dex-grown MP::MPΔPB1:GR and MP::MPΔPB1:GR was significant at *P*<0.05 (*) or *P*<0.001 (***) by *F*-test and *t*-test with Bonferroni correction. Sample population sizes: 26 (WT), 27 (*bdl*), 27 (MP::VP16:bdlΔI;*iaa12-1*;*tpl-1*), 18 (MP::MPΔPB1:GR), or 19 (dex-grown MP::MPΔPB1:GR) leaves; 265 (WT), 199 (*bdl*), 338 (MP::VP16:bdlΔI;*iaa12-1*;*tpl-1*), 248 (MP::MPΔPB1:GR), or 269 (dex-grown MP::MPΔPB1:GR) vein cell nuclei, and 316 (MP::VP16:bdlΔI;*iaa12-1*;*tpl-1*) flanking cell nuclei. (H-L) Dark-field illumination of cleared first leaves 14 DAG. (M) Cardinality index, connectivity index, and continuity index (mean ± SE) as defined in (Verna et al., 2015) and Materials & Methods; cyclicity index (mean ± SE) as defined in Materials & Methods. Difference between *bdl* and WT cardinality indices, between *bdl* and WT continuity indices, between *bdl* and WT connectivity indices, and between *bdl* and WT cyclicity indices, was significant at *P*<0.001 (***) by *F*-test and *t*-test with Bonferroni correction. Sample population sizes: WT, 30; *bdl*, 65; MP::VP16:bdlΔI;*iaa12-1*;*tpl-1*, 22; MP::MPΔPB1:GR, 42; dex-grown MP::MPΔPB1:GR, 38. Scale bars: (A-E) 25 µm; (I) 0.25 mm; (J) 0.5 mm; (H,K,L) 1 mm.

Were repression of *ATHB8* preprocambial expression by MP mediated by BDL, reducing or eliminating inhibition of MP transcriptional activity by BDL would lead to higher levels of *ATHB8* preprocambial expression in both vein and flanking cells, leading to broader *ATHB8* expression domains. To test this prediction, we turned the unstable BDL transcriptional repressor into a stabilized transcriptional activator as previously done for other IAA proteins (Tiwari et al., 2001; Tiwari et al., 2003; Li et al., 2009): we replaced the repressor domain of BDL (Li et al., 2011) with the activator domain of the *Herpes simplex* Virus Protein 16 (VP16) (Sadowski et al., 1988) and introduced a mutation that lengthens the half-life of BDL (Hamann et al., 2002). We expressed the resulting VP16:bdlΔI by the *ATHB8* promoter in the *iaa12-1*;*tpl-1* double mutant, which lacks *BDL* function (Overvoorde et al., 2005) and partially lacks the co-repressor function that mediates the IAA-protein-dependent repression of MP (Szemenyei et al., 2008). We quantified ATHB8::nYFP expression levels in second loops of 4-DAG first leaves of the resulting MP::VP16:bdlΔI;*iaa12-1*;*tpl-1* background.

Like in MP::MP, in MP::VP16:bdlΔI;*iaa12-1*;*tpl-1* ATHB8::nYFP expression levels were higher in flanking cells (Fig. 3A,D,F; Fig. 4A,C,G). Unlike in MP::MP, however, in MP::VP16:bdlΔI;*iaa12-1*;*tpl-1* ATHB8::nYFP expression levels were also higher in vein cells (Fig. 3A,D,F; Fig. 4A,C,G). Accordingly, stronger *ATHB8* expression domains in MP::VP16:bdlΔI;*iaa12-1*;*tpl-1* developing leaves were associated with stronger — though qualitatively similar — vein network defects in MP::VP16:bdlΔI;*iaa12-1*;*tpl-1* mature leaves: in the middle of these leaves, veins ran parallel to one another for the entire length of the narrow leaf laminae to give rise to wide midveins; toward the margin, veins ran close to one another for varying stretches of the laminae, then diverged, and either ran close to other veins or converged back to give rise to elongated meshes (Fig. 3G,J,K; Fig. 4H,J).

Next, we further tested the prediction that reducing or eliminating inhibition of MP transcriptional activity by BDL would lead to higher levels of *ATHB8* preprocambial expression in both vein and flanking cells, leading to broader *ATHB8* expression domains. As previously done (Krogan et al., 2012; Smetana et al., 2019; Amalraj et al., 2020), we created an irrepressible version of MP by deleting its PHOX/BEM1 (PB1) domain, which is required for IAA-protein-mediated repression (Tiwari et al., 2003; Wang et al., 2005; Krogan et al., 2012; Korasick et al., 2014). We fused the resulting MPΔPB1 to a fragment of the rat glucocorticoid receptor (GR) (Picard et al., 1988) to confer dexamethsone (dex)-inducibility, expressed the resulting MPΔPB1:GR by the *MP* promoter, and quantified ATHB8::nYFP expression levels in 4-DAG first leaves of the dex-grown MP::MPΔPB1:GR background.

Consistent with previous observations (Garrett et al., 2012; Krogan et al., 2012), in dex-grown MP::MPΔPB1:GR ATHB8::nYFP expression was no longer restricted to narrow stripes; instead, ATHB8::nYFP was expressed at higher levels in broad bands than spanned almost the entire width of the leaves (Fig. 4D,E,G). Accordingly, broader and stronger *ATHB8* expression domains in dex-grown MP::MPΔPB1:GR developing leaves were associated with veins running parallel to one another for the entire length of the narrow leaf laminae to give rise to midveins that spanned almost the entire width of dex-grown MP::MPΔPB1:GR mature leaves (Fig. 4H,K,L).

In conclusion, our results are consistent with the hypothesis that *MP* expression levels above a maximum threshold both activate and repress *ATHB8* preprocambial expression and that such repression of *ATHB8* preprocambial expression by MP is mediated by BDL.

### Relation Between *ATHB8* Expression Domains and Auxin Levels

IAA proteins, including BDL, are degraded in response to auxin (Gray et al., 2001; Tiwari et al., 2001; Zenser et al., 2001; Dharmasiri et al., 2005). Auxin-dependent degradation of BDL and other IAA proteins releases MP from inhibition, thus allowing MP to activate expression of its targets, including *BDL* and *ATHB8* (Hardtke et al., 2004; Weijers et al., 2005; Weijers et al., 2006; Donner et al., 2009; Ploense et al., 2009; Schlereth et al., 2010; Lau et al., 2011; Garrett et al., 2012; Krogan et al., 2012; Krogan et al., 2014; Wu et al., 2015). Therefore, narrow stripes of *ATHB8* preprocambial expression should correspond to peak levels of auxin. To test this prediction, we simultaneously imaged in midvein, first loops, and second loops of developing first leaves expression of ATHB8::nQFP (nuclear Turquoise Fluorescent Protein expressed by the *ATHB8* promoter) and of the auxin ratiometric reporter R2D2 (Liao et al., 2015), which expresses an auxin-degradable nYFP and a non-auxin-degradable nRFP by the *RIBOSOMAL PROTEIN S5A* promoter, which is highly active in developing leaves (Weijers et al., 2001). In the R2D2 reporter, a high RFP/YFP ratio thus indicates high levels of auxin, whereas a low RFP/YFP ratio indicates low levels of auxin (Liao et al., 2015).

At all tested stages, the RFP/YFP ratio was higher in ATHB8::nQFP-expressing cells than in cells flanking ATHB8::nQFP-expressing cells (Figure 5), suggesting that — consistent with previous observations (Mattsson et al., 2003; Scarpella et al., 2004) —domains of *ATHB8* preprocambial expression correspond to peak levels of auxin.

**Figure 5.**
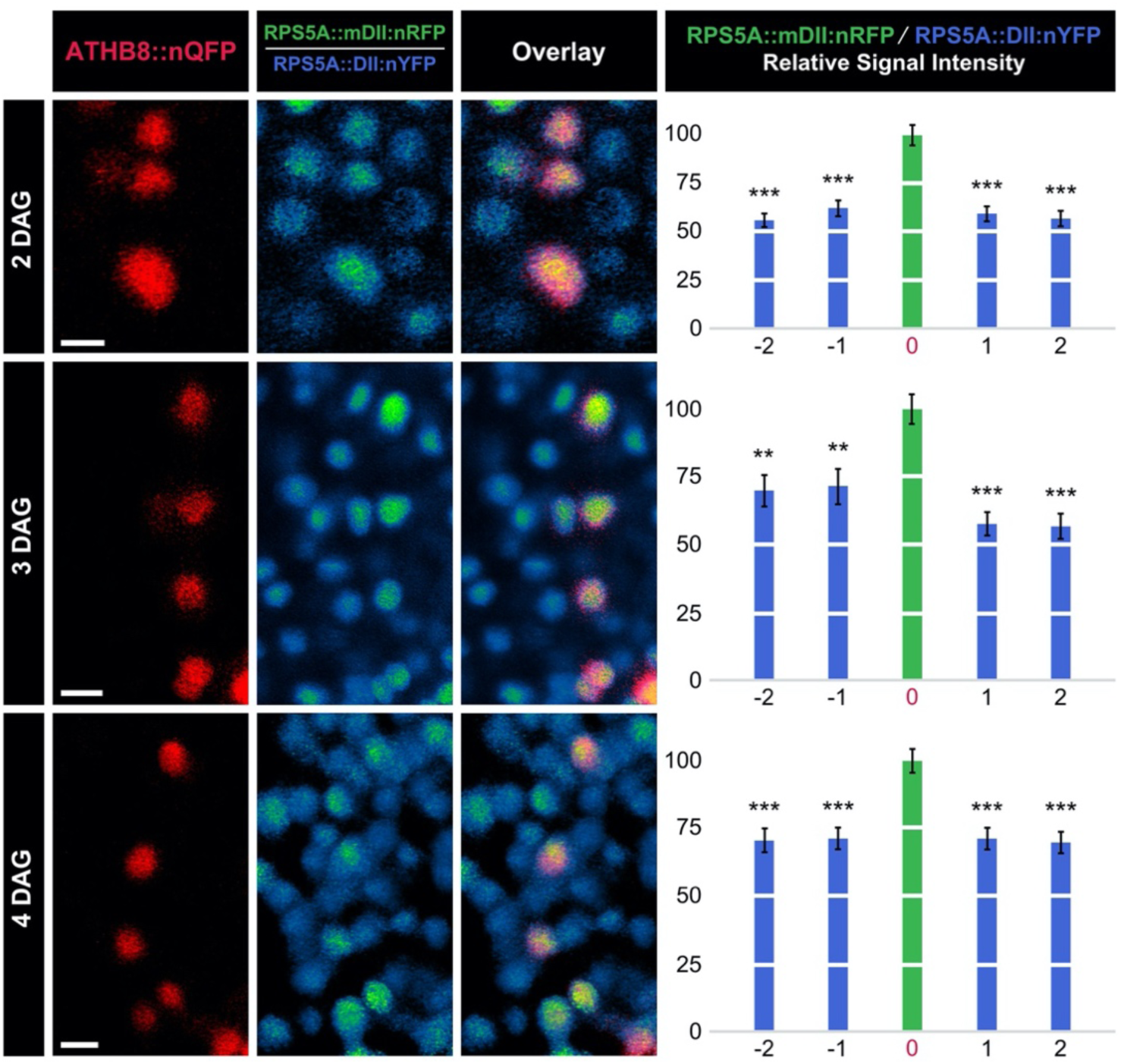
*ATHB8* Expression Domains and Auxin Levels. First leaves 2, 3, and 4 DAG. Columns 1–3: confocal laser scanning microscopy. Column 1: ATHB8::nQFP expression (red) associated with formation of midvein (2 DAG), first loop (3 DAG), or second loop (4 DAG) (Donner et al., 2009; Gardiner et al., 2011; Donner and Scarpella, 2013). Column 2: Ratio of RPS5A::mDII:nRFP expression to RPS5A::DII:nYFP expression. Look-up table visualizes expression ratio levels: high RPS5A::mDII:nRFP/RPS5A::DII:nYFP ratio (green) indicates high auxin levels; low RPS5A::mDII:nRFP/RPS5A::DII:nYFP ratio (blue) indicates low auxin levels. Column 3: overlays of images in columns 1 and 2; blue: low RPS5A::mDII:nRFP/RPS5A::DII:nYFP ratio, i.e. low auxin levels; yellow: co-expression of ATHB8::nQFP (red) and high RPS5A::mDII:nRFP/RPS5A::DII:nYFP ratio (green), i.e. high auxin levels. Column 4: Ratio of RPS5A::mDII:nRFP expression levels to RPS5A::DII:nYFP expression levels (mean ± SE) in nuclei flanking ATHB8::nQFP-expressing nuclei (positions “-2”, “-1”, “1”, and “2”) relative to ratio of RPS5A::mDII:nRFP expression levels to RPS5A::DII:nYFP expression levels in nuclei co-expressing ATHB8::nQFP (position “o”) during formation of midvein (top), first loop (middle), or second loop (bottom). Difference between ratio of RPS5A::mDII:nRFP expression levels to RPS5A::DII:nYFP expression levels in nuclei at position −2, −1, 1, or 2 and ratio of RPS5A::mDII:nRFP expression levels to RPS5A::DII:nYFP expression levels in nuclei at position 0 was significant at *P*<0.01 (**) or *P*<0.001 (***) by One-Way ANOVA and Tukey’s Pairwise test. Sample population sizes: 26 (2 DAG), 27 (3 DAG), or 29 (4 DAG) leaves; position −2: 56 (2 DAG), 42 (3 DAG), or 60 (4 DAG) nuclei; position −1: 52 (2 DAG), 37 (3 DAG), or 58 (4 DAG) nuclei; position 0: 74 (2 DAG), 85 (3 DAG), or 102 (4 DAG) nuclei; position 1: 44 (2 DAG), 44 (3 DAG), or 62 (4 DAG) nuclei; position 2: 42 (2 DAG), 25 (3 DAG), or 44 (4 DAG) nuclei. Scale bars (shown, for simplicity, only in column 2): 5 µm.

### Response of *ATHB8* Expression to Manipulation of MP-Binding Site Affinity

The hypothesis that *MP* expression levels below a minimum threshold are unable to activate *ATHB8* preprocambial expression predicts that reducing the affinity of MP for its binding site in the *ATHB8* promoter will lead to to extremely weak, or altogether absent, *ATHB8* preprocambial expression.

To test this prediction, we mutated the MP-binding site in the *ATHB8* promoter (TGTCTG) to lower (TGTCAG) or negligible (TAGCTG) affinity for MP-binding (Ulmasov et al., 1997; Ulmasov et al., 1999; Donner et al., 2009; Boer et al., 2014), and imaged nYFP expressed by the native or mutant promoters in second loops of 4-DAG first leaves.

Mutation of the MP-binding site in the *ATHB8* promoter to negligible affinity for MP-binding led to greatly reduced levels of nYFP expression (Fig. 6A,B,F), resembling near-complete loss of ATHB8::nYFP preprocambial expression in *mp-U55* (Donner et al., 2009) (Fig. 3A,B,F). Mutation of the MP-binding site in the *ATHB8* promoter to lower affinity for MP-binding led to lower levels of nYFP expression (Fig. 6A,C,F). Furthermore, expression along the domains was more heterogeneous than when nYFP was expressed by the native promoter (Fig. 6A,C,F), leading to seemingly fragmented domains of weak nYFP expression similar to those in *mp-11* (Fig. 3A,C,F) and *bdl* (Fig. 4A,B,G).

**Figure 6.**
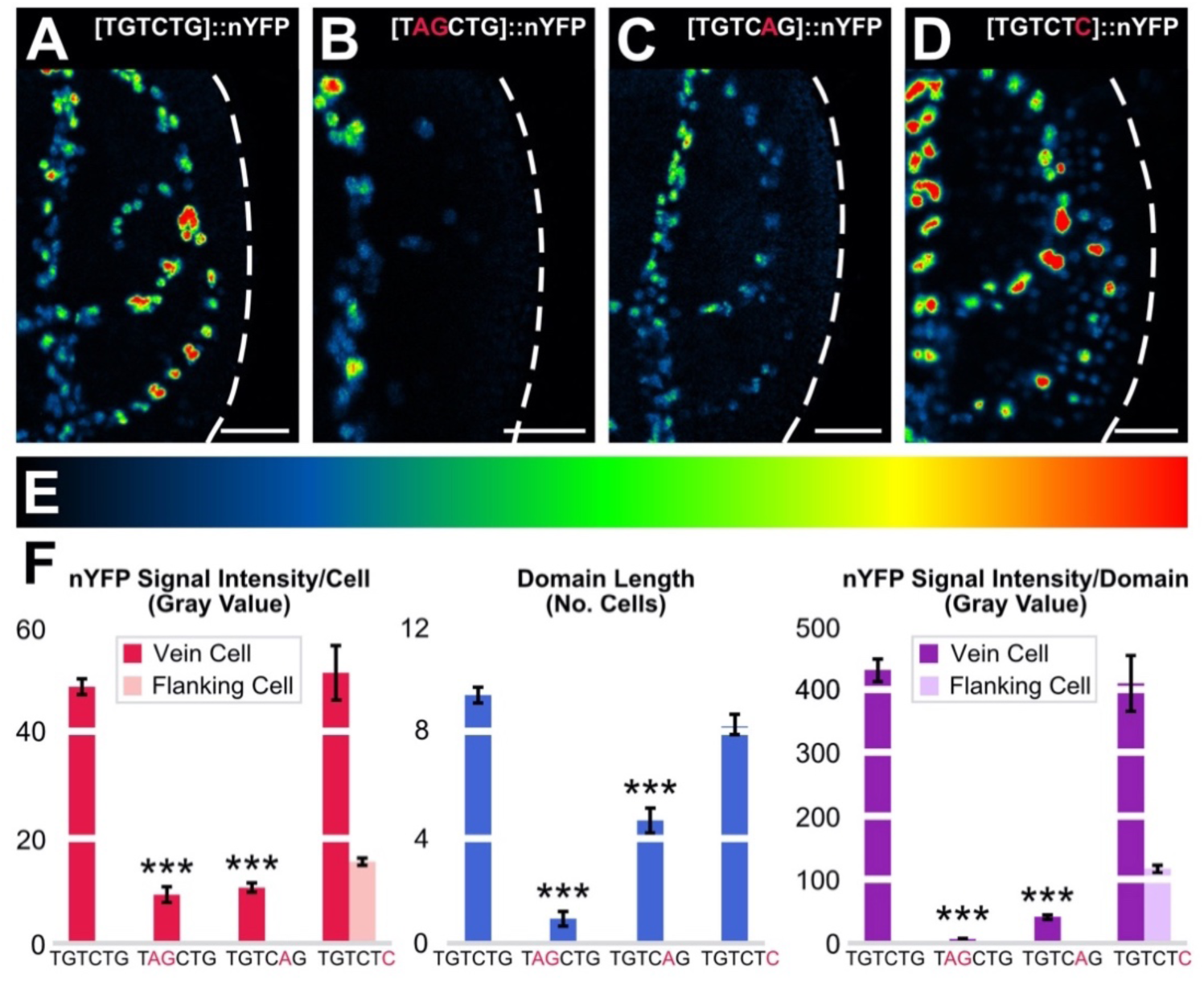
Activity of *ATHB8* Promoter Variants. (A–D) First leaves 4 DAG; confocal laser scanning microscopy; nYFP expression (look-up table — ramp in E — visualizes expression levels) driven by promoter variants (top right) with native ([TGTCTG]::nYFP≡ATHB8::nYFP) (A), negligible ([TAGCTG]::nYFP) (B), lower ([TGTCAG]::nYFP) (C), or higher ([TGTCTC]::nYFP) (D) affinity for MP-binding. Dashed white line: leaf outline. (F) nYFP expression level per cell expressed as mean gray value ± SE, nYFP expression domain length expressed as mean number of cells ± SE, and nYFP expression level per domain expressed as mean gray value ± SE. Difference between [TAGCTG]::nYFP and ([TGTCTG]::nYFP, and between [TGTCAG]::nYFP and ([TGTCTG]::nYFP was significant at *P*<0.001 (***) by *F*-test and *t*-test with Bonferroni correction. Sample population sizes: 21 ([TGTCTG]::nYFP), 22 ([TAGCTG]::nYFP), 21 ([TGTCAG]::nYFP), or 16 ([TGTCTC]:nYFP) leaves; 391 ([TGTCTG]::nYFP), 41 ([TAGCTG]::nYFP), 194 ([TGTCAG]::nYFP), or 261 ([TGTCTC]::nYFP) vein cell nuclei, and 611 ([TGTCTC]::nYFP) flanking cell nuclei. Scale bars: 25 µm.

The hypothesis that *MP* expression levels above a maximum threshold both activate and repress *ATHB8* preprocambial expression predicts that increasing the affinity of MP for its binding site in the *ATHB8* promoter will lead to higher levels of *ATHB8* preprocambial expression in flanking cells, leading to broader *ATHB8* expression domains, and to levels of *ATHB8* preprocambial expression in vein cells that are no lower — though not necessarily any higher — than those in WT.

To test this prediction, we mutated the MP-binding site in the *ATHB8* promoter (TGTCTG) to higher (TGTCTC) affinity for MP-binding (Ulmasov et al., 1997; Ulmasov et al., 1999; Donner et al., 2009), and imaged nYFP expressed by the native or mutant promoter in second loops of 4-DAG first leaves.

Mutation of the MP-binding site in the *ATHB8* promoter to higher affinity for MP-binding led to higher levels of nYFP expression in flanking cells (Fig. 6A,D,F), resulting in broad bands of nYFP expression similar to those in MP::MP (Fig. 3A,D,F) and, to a lesser extent, MP::VP16:bdlΔI;*iaa12-1*;*tpl-1* (Fig. 4A,C,G) and dex-grown MP::MPΔPB1:GR (Fig. 4D,E,G). However, unlike in MP::MP — in which ATHB8::nYFP expression levels in vein cells were lower than in WT (Fig. 3A,D,F) — and MP::VP16:bdlΔI;*iaa12-1*;*tpl-1* and dex-grown MP::MPΔPB1:GR — in which those levels were higher (Fig. 4A,C-E,G) — nYFP expression levels in vein cells were unchanged by mutation of the MP-binding site in the *ATHB8* promoter to higher affinity for MP-binding (Fig. 6A,D,F), suggesting that MP levels are normally nonlimiting for *ATHB8* preprocambial expression.

In conclusion, our results are consistent with the hypothesis that *MP* expression levels below a minimum threshold are unable to activate *ATHB8* preprocambial expression and that *MP* expression levels above a maximum threshold both activate and repress *ATHB8* preprocambial expression.

### An Incoherent Feedforward Loop Activating Gene Expression in Narrow Stripes

Consistent with interpretation of similar findings in animals (e.g., (Bellusci et al., 1997; Latinkic et al., 1997; Sato and Saigo, 2000)), our results suggest that an incoherent type-I feedforward loop (Mangan and Alon, 2003) restricts activation of expression of the plant gene *ATHB8* in preprocambial stripes and leads to vein network formation (Figure S5). Auxin activates *MP*, which in turn activates expression of intermediate-loop *AUX*/*IAA* genes like *BDL*. Both *MP* and *AUX*/*IAA* genes jointly regulate expression of *ATHB8*, which converts the auxin signal input into vein-network formation output. How *ATHB8* controls vein formation is unclear, but delayed procambium formation in *athb8* leaves (Donner et al., 2009) suggests that *ATHB8* promotes timely procambium formation, thereby preventing premature termination of initiation vein formation by mesophyll differentiation (Scarpella et al., 2004). Furthermore, because *athb8* enhances the defects in coordination of cell polarity and vein patterning induced by the inhibition of the polar, cell-to-cell transport of auxin (Donner et al., 2009), it is possible that *ATHB8* belongs to that auxin signaling pathway that controls coordination of cell polarity and vein patterning redundantly with polar auxin transport (Verna et al., 2019). However, this possibility remains to be tested.

In the future, it will be interesting to understand what generates peaks of auxin and *MP* levels at sites of vein formation. For example, like the *ATHB8*-related *PHABULOSA* in the root (Muller et al., 2016), *ATHB8* could control *MP* expression, such that interpretation of positional information fed back on generation of that information, as it often happens in animals (reviewed in (Jaeger et al., 2008)). Broader expression domains of an *MP* expression reporter in *athb8* leaves (Gagne et al., 2008; Donner et al., 2009) are consistent with such a possibility. Irrespective of how peaks of auxin and *MP* levels are generated, however, already now our results suggest a mechanism by which in plants a broadly expressed transcription factor activates target gene expression in narrow stripes. The very same regulatory mechanism that controls activation of *ATHB8* expression in single files of preprocambial cells is most frequently used in animals to generate stripes of gene expression (Cotterell and Sharpe, 2010), suggesting unexpected conservation of regulatory logic of striped gene-expression in plants and animals in spite of the independent evolution of their multicellularity. Nevertheless, in animals such regulatory logic typically leads to activation of target gene expression in a stripe that is outside the expression domain of the activating transcription factor (e.g., (Bellusci et al., 1997; Latinkic et al., 1997; Sato and Saigo, 2000; Yakoby et al., 2008)), whereas *ATHB8* expression is activated in a stripe that is a subset of the MP expression domain. It will be interesting to understand whether these are plant- and animal-specific outputs of the same conserved regulatory logic.

## MATERIALS & METHODS

### Plants

Origin and nature of lines, genotyping strategies, and oligonucleotide sequences are in Tables S1, S2, and S3, respectively. Seeds were sterilized and sowed as in (Sawchuk et al., 2008). Stratified seeds were germinated and seedlings were grown at 22°C under continuous light (∼90 µmol m^−2^ s^−1^). Plants were grown at 25°C under fluorescent light (∼100 µmol m^−2^ s^−1^) in a 16-h-light/8-h-dark cycle and transformed as in (Sawchuk et al., 2008).

### RT-qPCR

Total RNA was extracted with Qiagen’s RNeasy Plant Mini Kit from 4-day-old seedlings grown in half-strength Murashige and Skoog salts, 15 g l^−1^ sucrose, 0.5 g l^−1^ MES, pH 5.7, at 23°C under continuous light (∼80 µmol m^−2^ s^−1^) on a rotary shaker at 50 rpm. DNA was removed with Invitrogen’s TURBO DNA-free kit, and RNA was stabilized by the addition of 20 U of Thermo Fisher Scientific’s Superase-In RNase Inhibitor. First-strand cDNA was synthesized from ∼100 ng of DNase-treated RNA with Thermo Fisher Scientific’s RevertAid Reverse Transcriptase according to manufacturer’s instructions, except that 50 pmol of Thermo Fisher Scientific’s Oligo(dT)_18_ Primer, 50 pmol of Thermo Fisher Scientific’s Random Hexamer Primer, and 20 U of Superase-In RNase Inhibitor were used. qPCR was performed with Applied Biosystems’ 7500 Fast Real-Time PCR System on 2 µl of 1:3-diluted cDNA with 5 pmol of each gene-specific primers (Table S3), 2.5 pmol of gene-specific probe (Table S3), and Applied Biosystems’ TaqMan 2× Universal PCR Master Mix in a 10-µl reaction volume. Probe and primers were designed with Applied Biosystems’ Primer Express. Relative *MP* transcript levels were calculated with the 2^−Δ ΔCt^ method (Livak and Schmittgen, 2001) using *ACTIN2* transcript levels for normalization.

### Imaging

Developing leaves were mounted and imaged as in (Sawchuk et al., 2013), except that emission was collected from ∼1.5–5.0-µm-thick optical slices. In single-fluorophore marker lines, YFP was excited with the 514-nm line of a 30-mW Ar laser, and emission was collected with a BP 520–555 filter. In multiple-fluorophore marker lines, CFP, QFP, and autofluorescent compounds were excited with the 458-nm line of a 30-mW Ar laser; YFP was excited with the 514-nm line of a 30-mW Ar laser; and RFP was excited with the 543-nm line of a HeNe laser; CFP/QFP emission was collected with a BP 475–525 filter; YFP emission was collected with a BP 520–555 filter; RFP emission was collected between 581 and 657 nm; and autofluorescence was collected between 604 and 700 nm. Signal intensity levels of 8-bit grayscale images acquired at identical settings were quantified in the Fiji distribution of ImageJ (Schindelin et al., 2012; Schneider et al., 2012; Schindelin et al., 2015; Rueden et al., 2017). To visualize RFP/YFP ratios, the histogram of the YFP images was linearly stretched in the Fiji distribution of imageJ such that the maximum gray value of the YFP images matched that of the corresponding RFP images, and the RFP images were divided by the corresponding YFP images. Mature leaves were fixed, cleared, and mounted as in (Verna et al., 2019; Amalraj et al., 2020), and mounted leaves were imaged as in (Odat et al., 2014). Image brightness and contrast were adjusted by linear stretching of the histogram in in the Fiji distribution of ImageJ.

### Vein Network Analysis

The cardinality, continuity, and connectivity indices were calculated as in (Verna et al., 2015). Briefly, number of “touch points” (TPs, where a TP is the point where a vein end contacts another vein or a vein fragment), “end points” (EPs, where an EP is the point where an “open” vein — a vein that contacts another vein only at one end — terminates free of contact with another vein or a vein fragment), “break points” (KPs, where a KP is each of the two points where a vein fragment terminates free of contact with veins or other vein fragments), and “exit points” (XPs, where an XP is the point where a vein exits leaf blade and enters leaf petiole) in dark-field images of cleared mature leaves was calculated with the Cell Counter plugin in the Fiji distribution of ImageJ.

Because a vein network can be understood as an undirected graph in which TPs, EPs, KPs, and XPs are vertices, and veins and vein fragments are edges, and because each vein is incident to two TPs, a TP and an XP, a TP and an EP, or an XP and an EP, the cardinality index — a measure of the size (i.e. the number of edges) of a graph — is a proxy for the number of veins and is calculated as [(TPs+XPs-EPs)/2]+EPs, or (TPs+XPs+EPs)/2.

The continuity index quantifies how close a vein network is to a network with the same number of veins, but in which at least one end of each vein fragment contacts a vein, and is therefore calculated as the ratio of the cardinality index of the first network to the cardinality index of the second network: [(TP+XP+EP)/2]/[(TP+XP+EP+KP)/2], or (TP+XP+EP)/(TP+XP+EP+KP).

The connectivity index quantifies how close a vein network is to a network with the same number of veins, but in which both ends of each vein or vein fragment contact other veins, and is therefore calculated as the ratio of the number of “closed” veins — those veins which contact vein fragments or other veins at both ends — in the first network to the number of closed veins in the second network (i.e. the cardinality index of the second network): [(TP+XP-EP)/2]/[(TP+XP+EP+KP)/2], or (TP+XP-EP)/(TP+XP+EP+KP).

Finally, because the number of meshes in a vein network equals the number of closed veins, the cyclicity index — a proxy for the number of meshes in a vein network — is calculated as: (TP+XP-EP)/2.

## ACKNOWLEDGEMENTS

We thank the Arabidopsis Biological Resource Center for seeds of *athb8-27, iaa12-1*, and *tpl-1*; Dolf Weijers for seeds of *mp-B4149* and R2D2, Gerd Jurgens for *bdl* seeds, Hiroo Fukuda and Kyoko Ohashi-Ito for SHR::mATHB8 DNA, and Zachary Nimchuk for nQFP DNA. We thank Neil Harris and Przemek Prusinkiewicz for helpful comments on the manuscript. This work was supported by Discovery Grants of the Natural Sciences and Engineering Research Council of Canada (NSERC) (RGPIN-2016-04736 to E.S.). A.K. was supported, in part, by an NSERC CGS-M Scholarship. J.L.G. was supported, in part, by an NSERC CGS-M scholarship and an NSERC PGS-D Scholarship. T.J.D. was supported, in part, by an NSERC CGS-D Scholarship and an Alberta Ingenuity Student Scholarship.

