## Supplemental Material for "Control of Vein-Forming, Striped Gene-Expression by Auxin Signaling"

### SUPPLEMENTAL TABLES

*Table S1. Origin and Nature of Lines*

| LINE | ORIGIN / NATURE |
|---|---|
| *athb8-11* | ABRC (CS6969); (Prigge et al., 2005); WT at the *ER* (AT2G26330) locus |
| *athb8-27* | ABRC (CS111153) |
| SHR::miR165a | Transcriptional fusion of *SHR* (AT4G37650; -2505 to -10; primers: "SHR HindIII F" and "SHR SalI R") to miR165a (AT1G01183; - 138 to +323 relative to the transcriptional start-site; primers: "SalI FWD – MiRNA 165" and "KpnI REV – MiRNA 165") |
| SHR::mATHB8 | (Ohashi-Ito et al., 2013) |
| SHR::mATHB8:EAR | Translational fusion of SHR::mATHB8 (Ohashi-Ito et al., 2013) (primers: "SalI SHR Promoter FP" and "XhoI mATHB8 RP") to the sequence encoding the EAR portable repressor domain (Hiratsu et al., 2003) (primers: "EAR XhoI + KpnI Forward" and "EAR Reverse") |
| MP::ATHB8 | Transcriptional fusion of *MP* (AT1G19850; -3281 to -1; primers: "MP BamHI Fwd" and "MP KpnI Rev") to the *ATHB8* (AT4G32880) cDNA (GeneBank accession: BT008798; ABRC: U24724; |

| | |
|---|---|
| | +1 to +2502; primers: "ATHB8 cDNA KpnI FWD" and "ATHB8 cDNA SmaI Rev") |
| MP::mATHB8 | Transcriptional fusion of *MP* (AT1G19850; -3281 to -1; primers: 63 "MP BamHI Fwd" and "MP KpnI Rev") to the *ATHB8* (AT4G32880) cDNA (GeneBank accession: BT008798; ABRC: U24724; +1 to +2502; primers: "ATHB8 cDNA KpnI FWD" and "ATHB8 cDNA SmaI Rev"; "ATHB8mut165FWD" and "ATHB8mut165REV") |
| ATHB8::nCFP | (Sawchuk et al., 2007) |
| MP::MP:YFP | Translational fusion of *MP* (AT1G19850; -3281 to +3815; primers: "MP Prom SalI Fwd" and "MP KpnI Rev-2"; "MP 3 kb SalI Fwd" and "MP 3 kb XhoI Rev") to the sequence encoding EYFP (primers: "ECFP AflII F" and "ECFP AflII R"); rescues the root (240/240 seedlings), vein (Figure S1), and inflorescence (160/160 plants) defects of *mp-B4149* |
| *mp-B4149* | (Weijers et al., 2005) |
| RIBO::nCFP | ABRC (CS23898); (Gordon et al., 2007); WT at the *ER* (AT2G26330) locus |
| ATHB8::nYFP | (Sawchuk et al., 2007) |
| *mp-U55* | ABRC (CS8147); (Mayer et al., 1993; Donner et al., 2009) |

| | |
|---|---|
| *mp-11* | (Odat et al., 2014) |
| MP::MP | *MP* (AT1G19850; -3281 to +3830; primers: "MP Prom SalI Fwd" and "MP KpnI Rev-2"; "MP 3KB SalI Fwd" and "MP 3kb XhoI Rev"); rescues the root (169/176 seedlings), vein (Figure S1), and inflorescence (6/6 plants) defects of *mp-B4149* |
| *bdl* | (Hamann et al., 1999); introgressed into Col-0 |
| MP::VP16:bdlΔI | Transcriptional fusion of *MP* (AT1G19850; -3281 to -1; primers: "MP BamHI Fwd" and "MP KpnI Rev-1") to a translational fusion of the sequence encoding the activation domain of the *Herpes simplex* virus protein 16 (VP16) (Sadowski et al., 1988) (primers: "VP16 NcoIF2" and "VP16 PstIR") to a 5′- terminally deleted *bdl* (Hamann et al., 2002) (+94 to +1229; primers: "BDL PstIF" and "BDL BamHIR"; "BDL mut F1", "BDL mut F2", "BDL mut F3", "BDL mut F4", "BDL PstIF", and "BDL MfeI mut R"; "BDLd1 PstI F" and "BDL BAMHI R") |
| *iaa12-1* | ABRC (CS25213); (Overvoorde et al., 2005) |
| *tpl-1* | ABRC (CS65909); (Long et al., 2002) |
| MP::MPΔPB1:GR | Translational fusion of *MP* (AT1G19850; -3427 to +2388; primers: "MP SalI Forward – Primer # 2" and "MP EcoRI Reverse") to the sequence encoding a fragment of the rat glucocorticoid receptor (GR) (Aoyama and Chua, 1997) |

| | |
|---|---|
| | (primers: "SpeI GR Forward" and "SacII + KpnI (Internal) GR Reverse") |
| ATHB8::nQFP | Transcriptional fusion of *ATHB8* (AT4G32880; -2070 to -1; primers: "SalI 2KB ATHB8 Promoter Forward" and "ApaI 2KB ATHB8 Promoter Reverse") to the sequence encoding 2xmTQ2-N7 (primers: "ApaI 2xmTurqoise Forward" and "KpnI 2xmTFP Reverse") |
| R2D2 | (Liao et al., 2015) |
| [TGTCTG]::nYFP | (Donner et al., 2009) |
| [TAGCTG]::nYFP | (Donner et al., 2009) |
| [TGTCAG]::nYFP | Transcriptional fusion of *ATHB8* (AT4G32880; -953 to -1; primers: "1NagARE" and "Athb8 R-5") to the sequence encoding HTA6:EYFP (Zhang et al., 2005) |
| [TGTCTG]::nYFP | Transcriptional fusion of *ATHB8* (AT4G32880; -953 to -1; primers: "1NcARE" and "Athb8 R-5") to the sequence encoding HTA6:EYFP (Zhang et al., 2005) |

*Table S2. Genotyping Strategies*

| LINE | STRATEGY |
| --- | --- |
| *athb8-11* | ATHB8: "Athb8 0.5" and "athb8attB2R"; *athb8-11*: "athb8 -5944" and "PD991- RB |
| *athb8-27* | ATHB8: "athb8-27 RP" and "athb8-27 LP"; *athb8-27*: "athb8-27 RP" and "Spm32" |
| *mp-B4149* | "MP 1498-s" and "MP2082-AS"; *Mse*I |
| *mp-U55* | "MP Seq 2061" and "U55 Geno Rev"; *Sml*I |
| *mp-11* | *MP*: "Sail_1265_F06LP" and "Sail_1265_F06RP"; *mp-11*: "LB3" and "Sail_1265_F06RP" |
| *bdl* | "bdl geno F" and "bdl geno R"; *Hae*III |
| *iaa12-1* | *IAA12*: "SALK_138684 LP" and "SALK_138684 RP"; *iaa12-1*: "LBb1.3" and "SALK_138684 RP" |
| *tpl-1* | "tpl Caps Genotyping Forward" and "tpl Caps Genotyping Reverse"; *Nco*I |

*Table S3. Oligonucleotide Sequences*

| NAME | SEQUENCE (5' TO 3') |
|---|---|
| SHR HindIII F | GAGAAGCTTGACAAAGAAGCAGAGCGTGG |
| SHR SalI R | TGGGTCGACTTAATGAATAAGAAAATGAATAGAAGAAAGGG |
| SalI FWD – MiRNA 165 | ATTGTCGACCCACTCATCATTCCCTCATC |
| KpnI REV – MiRNA 165 | AGCGGTACCCTTATAGAAAATACTTCGTTAGCTTG |
| SalI SHR Promoter FP | GGGGTCGACACATAAACCAGTAGACAT |
| XhoI mATHB8 RP | GGGCTCGAGTATAAAAGACCAGTTGAGG |
| EAR XhoI + KpnI Forward | TCGAGCTAGATCTGGATCTAGAACTCCGTTTGGGTTTCGCTTAAGGTAC |
| EAR Reverse | CTTAAGCGAAACCCAAACGGAGTTCTAGATCCAGATCATGC |
| MP BamHI Fwd | AAGGGATCCTCCGGGTTAATCAGTATTATTAC |
| MP KpnI Rev | ACAGGTACCACAGAGAGATTTTTCAATGTTCTG |
| ATHB8 cDNA KpnI FWD | GTCGGTACCATGGGAGGAGGAAGCAATAATAG |
| ATHB8 cDNA SmaI Rev | ATGCCCGGGATCATATAAAAGACCAGTTGAGG |
| ATHB8mut165FWD | ATAGGAATCGTTGCTATTTCTC |

| | |
|---|---|
| ATHB8mut165REV | GGAATCTGGTCCAGGCTTCATC |
| MP Prom SalI Fwd | CCCGTCGACGTATATATAAACAATACCACCTTATAAC |
| MP KpnI Rev-2 | CATGGTACCTGCAGAATTAGCATACCACAC |
| MP 3 kb SalI Fwd | TCTGTCGACTCCGGGTTAATCAGTATTATTAC |
| MP 3 kb XhoI Rev | ATTCTCGAGTTAAGAGTTAAGACCACCTCC |
| ECFP AflII F | TTACTTAAGGTGAGCAAGGGCGACGAGC |
| ECFP AflII R | AGACTTAAGATTGTACAGCTCGTCCATGCC |
| VP16 NcoIF2 | TTACCATGGCCCCCCCGACCGATGTC |
| VP16 PstIR | TTTCTGCAGCCCCACCGTACTCGTCAATTC |
| BDL PstIF | ATACTGCAGCTCGTGGTGTGTCAGAATTGGAC |
| BDL BamHIR | TACGGATCCACTAAACTGGGTTGTTTCTTTGTC |
| BDL mut F1 | AATCTTCCGGCGGAGAGTGTTAGAGAATTGGG |
| BDL mut F2 | GTGGGTAAAAGTAATCTTCCGGCGGAGAGTG |
| BDL mut F3 | GTGTCAGAATTGGAGGTGGGTAAAAGTAATCTTCCG |
| BDL mut F4 | CGTGGTGTGTCAGAATTGGAGGTGGGGAAGAGTAATC |
| BDL MfeI mut R | TAACAATTGGTGACCATCCTACCACTTGAC |
| BDLd1 PstI F | AAACTGCAGCGTGGAAAGAGCGTGGG |

| | |
|---|---|
| MP SalI Forward – Primer # 2 | GGGGTCGACCGGATTCGTGATCTTCGTATCCCAT |
| MP EcoRI Reverse | ATTGAATTCGGTTCGGACGCGGGGTGTCGCAATT |
| SpeI GR Forward | GGGACTAGTGGAGAAGCTCGAAAAACAAAG |
| SacII + KpnI (Internal) GR Reverse | AATCCGCGGGGTACCTCATTTTTGATGAAACAGAAG |
| SalI 2KB ATHB8 Promoter Forward | CGCGTCGACCATTATAAATATCACGACTGTA |
| ApaI 2KB ATHB8 Promoter Reverse | ATTGGGCCCCTTTGATCCTCTCCGATCTCT |
| ApaI 2xmTurqoise Forward | ATTGGGCCCATGGTGAGCAAGGGCGAGGA |
| KpnI 2xmTFP Reverse | CGAGGTACCTCACTCTTCTTCTTGATCAGCTTCTG |
| 1NagARE | GGGGACAAGTTTGTACAAAAAAGCAGGCTTGGTTGT CTCGTATTAAGGG |
| Athb8 R-5 | GGGGACCACTTTGTACAAGAAAGCTGGGTCTTTGAT CCTCTCCGATCTCTC |
| 1NcARE | GGGGACAAGTTTGTACAAAAAAGCAGGCTTGGTTAC CTGGTATTAAGGG |
| athb8-27 FP | TGTGAAGAATGGATCCACCTC |
| athb8-27 RP | AGTGGTCAACACCACTTGACC |
| Spm32 | TACGAATAAGAGCGTCCATTTTAGAGTG |

| | |
|---|---|
| Athb8 0.5 | GGGGACAAGTTTGTACAAAAAAGCAGGCTTCCTTTG CTTCCAGAGACCAGCG |
| athb8attB2R | GGGGACCACTTTGTACAAGAAAGCTGGGTCTTTGAT CCTCTCCGATCTCTC |
| athb8 -5944 | GGTTTGGCATAAAAGTGCGG |
| PD991- RB | AAAACCTGGCGTTACCCAACT |
| MP 1498-s | CTCTCAGCGGATAGTATGCACATCGG |
| MP2082-AS | ATGGATGGAGCTGACGTTTGAGTTC |
| MP Seq 2061 | CATAATGTTACTCTTCATGTACGCC |
| U55 Geno Rev | GTGCTGTTTGTTGGCGATTGG |
| Sail_1265_F06LP | GCTTCATCTCTTCAAGCAAGG |
| Sail_1265_F06RP | TCCCAAAGTCTCACCACTCAC |
| LB3 | TAGCATCTGAATTTCATAACCAATCTCGATACAC |
| bdl geno F | GCTCAAATCTTGTGATGTGAGTG |
| bdl geno R | AGTCCACTAGCTTCTGAGGTTCCC |
| SALK_138684 LP | GTGGGGAAGAGTAATCTTCCG |
| SALK_138684 RP | CTTCTGCTCTTGACGTCTTGG |
| LBb1.3 | ATTTTGCCGATTTCGGAAC |

| | |
|---|---|
| tpl Caps Genotyping Forward | GCCCTGAAAATGACATCGGT |
| MP PrimeTime Probe | /56-FAM/CAGACTCAC/ZEN/ AGGCCTTCTCTCGCCA/3IABKFQ/ |
| MP PrimeTime Primer 2 | TGTACCAGTGCCTCCAGAATTATC |
| MP PrimeTime Primer 1 | TCCAGTCGCAGATCACATCAG |
| ACT2 PrimeTime Probe | /56-FAM/ACAGCACTT/ZEN/ GCCCAAGAGCATGA/3IABKFQ/ |
| ACT2 PrimeTime Primer 2 | TACTTCCTTTCAGGTGGTGCA |
| ACT2 PrimeTime Primer 1 | GCTGACCGTATGAGCAAAGAAAT |

### SUPPLEMENTAL FIGURE LEGENDS

*Figure S1. MP::MP:YFP and MP::MP Functionalities in Vein Network Formation*

Dark-field illumination of cleared first leaves 14 DAG. Top right: genotype. Scale bars: 0.5 mm.

*Figure S2.* ATHB8 *Expression Domains and MP and* RIBO *Expression Levels*

First leaves 4 DAG. Confocal laser scanning microscopy. Top right: reporter. Dashed green outline: second loop nuclei expressing ATHB8::nCFP (A,B) or ATHB8::nYFP (D,E). (B,E) Look-up table — ramp in C — visualizes expression levels. Scale bars (shown, for simplicity, only in A and D): 5 μm.

*Figure S3.* ATHB8 *Expression Domains and* RIBO *Expression Levels*

(A–F) First leaves 4 DAG. (A) Schematic of 4-DAG leaf — imaged in B–E — illustrating onset of *ATHB8* expression (red) — imaged in B — associated with second loop formation (Donner et al., 2009; Gardiner et al., 2011; Donner and Scarpella, 2013); increasingly darker gray: progressively older *ATHB8* expression domains. (B–E) Confocal laser scanning microscopy. (B) ATHB8::nYFP expression. (C) RIBO::nCFP expression. (D) Autofluorescence. (E) Overlay of images in B–D; red: ATHB8::nYFP expression; green: RIBO::nCFP expression; blue: autofluorescence. (F) RIBO::nCFP expression levels (mean ± SE) in nuclei at positions -2, -1, 1, and 2 — as defined in legend to Figure 3 — relative to RIBO::nCFP expression levels in nuclei at position 0 — as defined in legend to Figure 3 — during second loop formation. Difference between RIBO::nCFP expression levels in nuclei at position -2 or -1 and RIBO::nCFP expression levels in nuclei at position 0 was significant at $P<0.001$ (***) by One-Way ANOVA and Tukey's Pairwise test. Sample population sizes: 27 leaves; position -2, 42 nuclei; position -1, 64 nuclei; position 0, 69 nuclei; position 1, 50 nuclei; position 2, 28 nuclei. Scale bars (shown, for simplicity, only in column 2): 5 μm.

*Figure S4.* mp-11 *and MP::MP Effects on* MP *Expression*

*MP* transcript levels in *mp-11* and MP::MP seedlings relative to *MP* transcript levels in WT (mean ± SE of three technical replicates for each of three biological replicates); seedlings 4 DAG; RT-qPCR. Difference between *mp-11* and WT, and between MP::MP and WT was significant at $P<0.001$ (***) by $F$-test and $t$-test with Bonferroni correction.

*Figure S5. Summary and Interpretation.*

A three-gene incoherent type-I feedforward loop (Mangan and Alon, 2003) activates *ATHB8* expression in narrow preprocambial stripes and leads to vein network formation. *MP* receives the auxin input and activates expression of intermediate-loop *AUX/IAA* genes like *BDL/IAA12*. Both *MP* and *AUX/IAA* genes jointly regulate expression of the stripe gene *ATHB8*, which converts the auxin input into vein-network formation output. Arrows indicate positive effects; blunt-ended lines indicate negative effects.

### SUPPLEMENTAL FIGURES

*Figure S1*

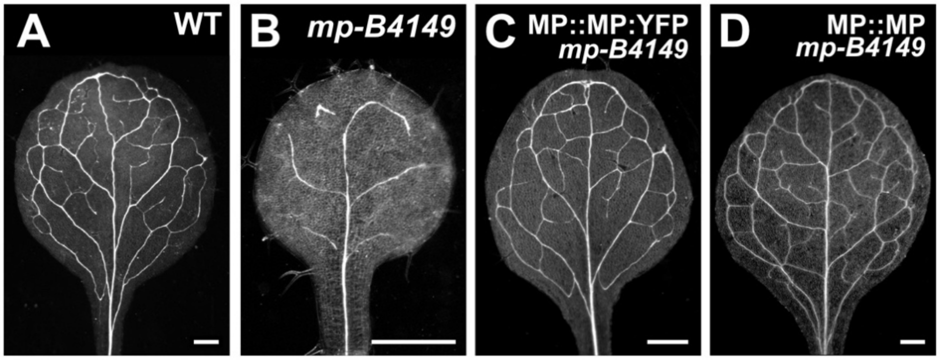

*Figure S2*

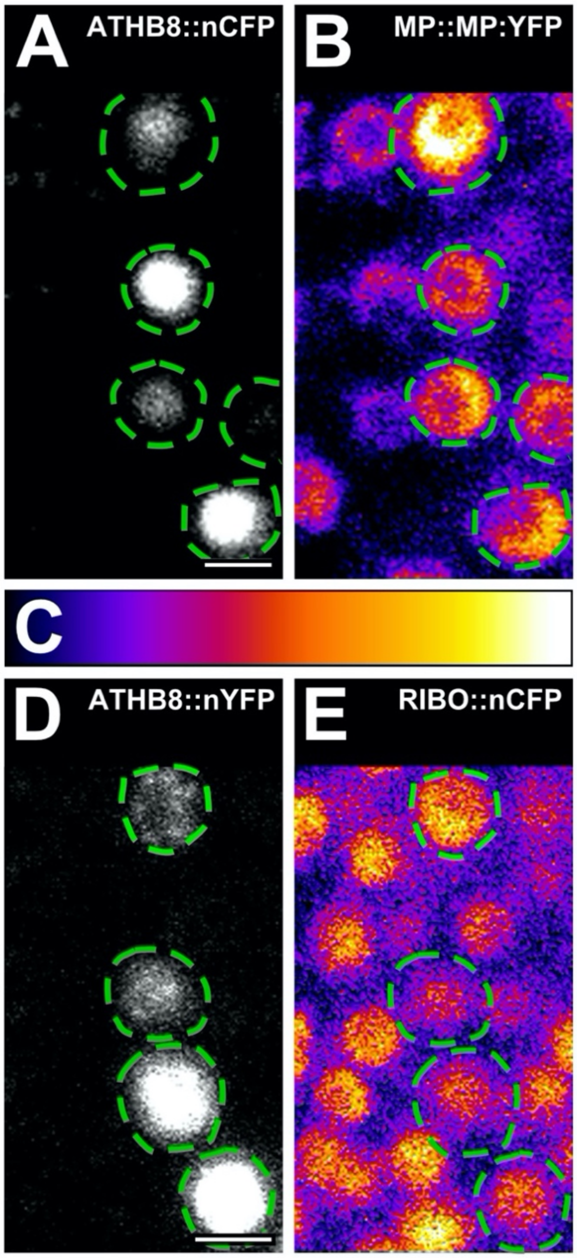

*Figure S3*

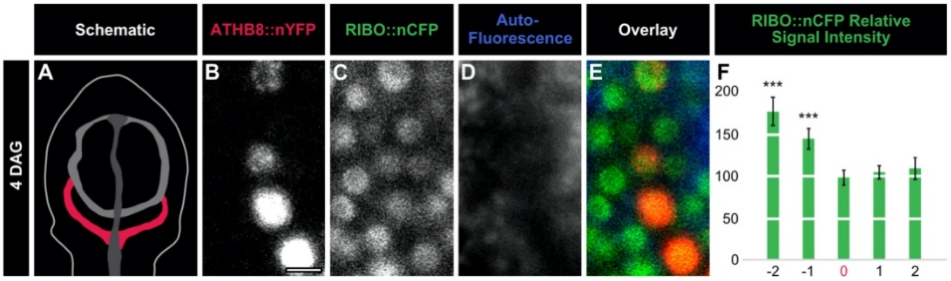
Schematic
ATHB8::nYFP
RIBO::nCFP
Auto-
Fluorescence
Overlay
RIBO::nCFP Relative
Signal Intensity
4 DAG
A
B
C
D
E
F
***
***
200
150
100
50
0
-2
-1
0
1
2

*Figure S4*

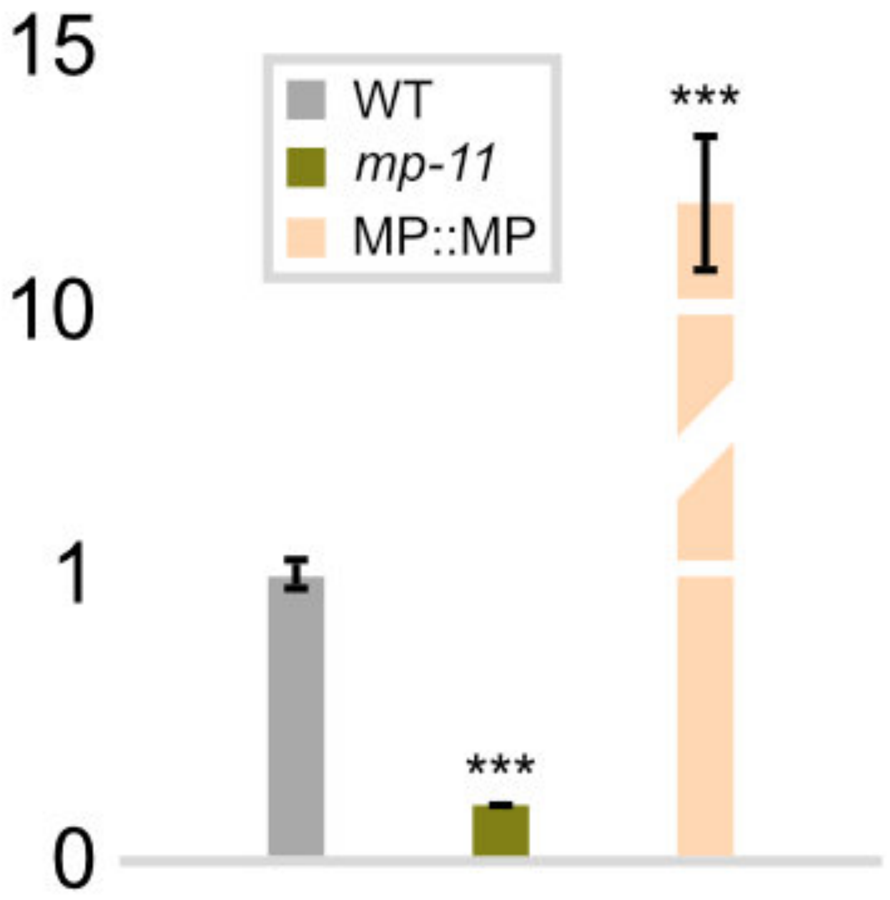

*Figure S5*

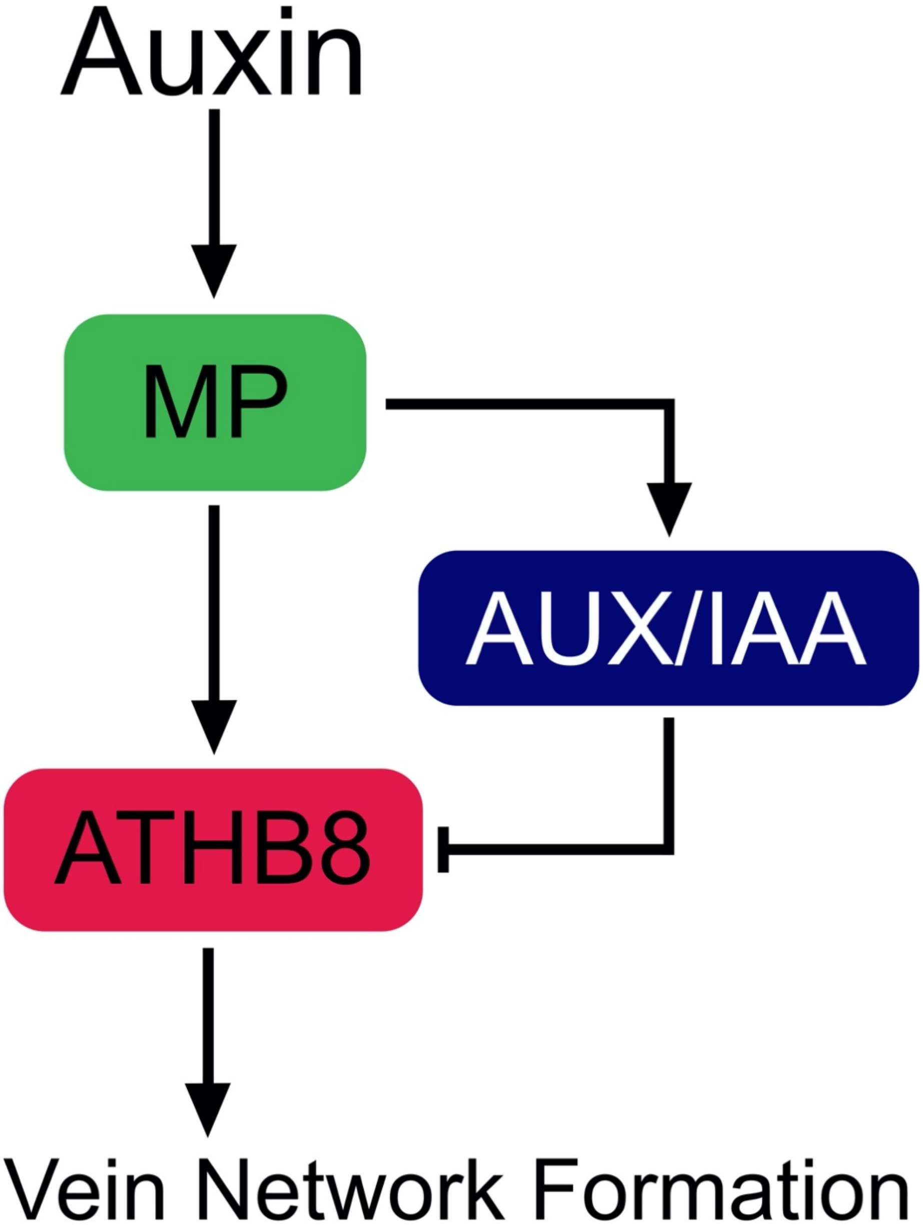
